# Deriving Accurate Lipid Classification based on Molecular Formula

**DOI:** 10.1101/572883

**Authors:** Joshua M. Mitchell, Hunter N.B. Moseley

## Abstract

**Introduction:** Although Fourier-transform mass spectrometry has substantially improved our ability to detect lipids and other metabolites; the untargeted and accurate assignment of detected metabolites remains an unsolved problem in metabolomics. New assignment methods such as our SMIRFE algorithm can assign elemental molecular formula to observed spectral features in an untargeted manner without orthogonal information from tandem MS or chromatography. However, for many lipidomics applications, it is necessary to know at least the lipid category or class that is associated with a detected spectral feature in order to derive biochemical interpretation.

**Objectives:** Our goal is to develop a method for robustly classifying elemental molecular formula assignments into lipid categories for application to SMIRFE-generated assignments.

**Results:** Using machine learning, we developed a method that can predict lipid category and class from SMIRFE molecular formula assignments. Our methods achieve high accuracy (>90%) and precision (>83%) for all eight of the lipid categories in the LIPIDMAPS database. Model performance was evaluated using sets of theoretical, data-derived, and artifactual molecular formulas. Our models were generalizable, applicable to real-world datasets, and very discriminating with most molecular formulas classified to the “not lipid” category. Lipid categories with the highest classification propensities were glycerophospholipids and sphingolipids, matching the highest category prevalence in LIPIDMAPS.

**Conclusions:** Our methods enable the lipid classification of untargeted molecular formula assignments generated by SMIRFE without orthogonal information, facilitating biochemical interpretation of highly untargeted lipidomics experiments. However, this lipid classification appears insufficient for validating single-spectrum assignments, but could be useful in cross-spectrum assignment validation.

## Introduction

Lipidomics is the subdiscipline of metabolomics concerned with the analytical investigation of the lipidome, the set of lipid metabolites and their roles within the metabolome. Unlike other categories of metabolites, that are largely grouped based on their structures, lipids are defined by their very low solubility in water and collectively represent a structurally and chemically diverse set of metabolites with various roles in normal and pathological cellular function. By virtue of this structural and chemical diversity, which often confers amphipathic properties, seemingly every life process involves lipids, including but not limited to: maintenance of cellular structure (Singer and Nicolson, 1972); membrane fluidity (Clamp *et al.*, 1997) (Chen and Yu, 1994); intracellular, extracellular, and hormonal signaling (Zechner *et al.*, 2012) (Morrison and Farmer, 2000); energy metabolism (J R Neely and Morgan, 1974) (Adeva-Andany *et al.*, 2018); and disease processes (De Pablo and De Cienfuegos, 2000) including cancer (Zhang and Du, 2012) (Ray and Roy, 2018). Thus, through lipidomics, more complete modeling of cellular metabolism and a better understanding of physiological and pathological processes at the mechanistic level can be achieved (Lydic and Goo, 2018).

Although the potential benefit of lipidomics are enormous, the rigorous analytical investigation of the lipidome in real-world biological samples requires the reliable observation of lipid features in the samples as well as the accurate assignment of those features to a lipid structure and/or lipid class. This represents a significant bioanalytical chemistry problem due to the high structural diversity of lipids, their wide range of observed concentrations, and differences in lipid profiles between compartments and with respect to time (Horvath and Daum, 2013) (Aviram *et al.*, 2016) (Fahy *et al.*, 2005). Given its sensitivity to a wide range of chemical structures and low detection limits, mass spectrometry remains the most popular analytical technique for lipidomics analysis (Köfeler *et al.*, 2012). Traditionally, mass spectrometry has been used in conjunction with other analytical techniques such as gas chromatography (Quehenberger *et al.*, 2011), liquid chromatography (Masoodi and Nicolaou, 2006), (Sandra *et al.*, 2010) or TLC (Valdes-Gonzalez *et al.*, 2011) to provide additional orthogonal information that aid in the assignment of observed lipid features. Recent advances in mass spectrometry, namely Fourier transform mass spectrometry (FT-MS), have provided significant improvements in mass accuracy, mass resolution, and sensitivity (Eliuk and Makarov, 2015). Together, these analytical improvements provide the capability to resolve distinct isotopologues with identical unit masses, which in turn enables natural abundance correction for multi-isotope labeling experiments (Carreer *et al.*, 2013; Moseley, 2010), improved assignment accuracy without orthogonal chemical information (Moseley *et al.*, 2018), and the detection of compounds in the sub-femtomolar range (Eyles and Kaltashov, 2004) (Dettmer *et al.*, 2007). These capabilities enable the use of stable isotope resolved metabolomics (SIRM) techniques in combination with traditional lipidomics methodologies (Li *et al.*, 2013) that can provide richer information, allowing the elucidation of unknow metabolic pathways, the quantification of relative fluxes through connected metabolic pathways, and the identification of active metabolic pathways under various cellular conditions (Postle and Hunt, 2009) (Allen *et al.*, 2015).

Although mass spectrometry, especially FT-MS, enables the robust *detection* of lipid features, the assignment of those features to lipids and by extension to lipid class, remains challenging. Although mass spectrometry enables the detection of previously unobserved lipids, existing spectral assignment methodologies such as LipidSearch (Peake *et al.*, 2013) and PREMISE (Lane *et al.*, 2009) rely heavily or exclusively on observed m/z values from MS1 to query databases of known metabolites for assignment. This can result in either a lack of assignments for these features or worse, incorrect assignments for these features which can cause large interpretive errors later in an analysis. The presence of spectral artifacts in FT-MS spectra can result in the consistent misassignment of artifactual features, leading to substantial errors in downstream analyses (Mitchell *et al.*, 2018). Additionally, the potential for assignment bias from using these databases significantly hampers the potential for discovery (Moseley, 2013) – a stated goal of many untargeted lipidomics analyses. Although orthogonal information from chromatography or MS/MS can cross validate potential assignments; the necessary combined analytical setups are more complex, generate additional information that must be processed, require larger amounts of sample (Chekmeneva *et al.*, 2017), are incompatible with direct infusion experiments, and can still suffer from assignment bias when using fragmentation or retention time annotated databases. The incompleteness of metabolite and lipid databases (Mitchell *et al.*, 2014) (Schrimpe-Rutledge *et al.*, 2016) is a major source of assignment error that cannot be easily overcome through additional orthogonal information.

Since orthogonal chromatographic information is not a panacea, advances in FT-MS assignment techniques and improvements in electrospray ionization have made direct-infusion mass spectrometry an increasingly popular analytical setup for both metabolomics and lipidomics. Using assignment techniques such as our in-house SMIRFE algorithm (Moseley *et al.*, 2018) (Mitchell *et al.*, 2019), elemental molecular formulae can be robustly assigned to observed spectral features without information from chromatography or MS/MS and without querying existing databases of metabolite and lipid structures. This assignment methodology is ideally suited for untargeted metabolomics and lipidomics workflows, with the molecular formula assignments useful for both biomarker characterization and relative metabolic flux analysis after natural abundance correction (Moseley, 2010) (Carreer *et al.*, 2013). Furthermore, this assignment methodology is resilient against misassignment due to common artifacts in FT-MS spectra. However, many lipidomics experiments are concerned with changes at the lipid class level, which neither SMIRFE molecular formula assignments nor elemental analysis by other methods like inductively coupled plasma mass spectrometry directly provide.

Lipid classes are sets of lipids that share certain chemical structure features. When the chemical structure of a lipid feature is known, either through database lookup or other analytical approaches, classification of that structure into a lipid class is straightforward. Automated tools such as ClassyFire (Djoumbou Feunang *et al.*, 2016) use machine learning methods to automatically assign lipid class (and more generally metabolite class) based on the input structures; however, structural information cannot be directly acquired through elemental formula assignment methods. Although detection of potential lipid features can be achieved using ratios of heteroatoms (Brockman *et al.*, 2018), the classification of molecular formulae into specific lipid classes remains an unsolved problem that can prevent the effective biochemical interpretation of SMIRFE-generated formulas derived from class-level lipidomics analyses.

Manually constructing rules that can map elemental formulae to lipid classes is a daunting proposition and would result in rules that are fragile and likely incomplete and incorrect. Fortunately, the prediction of lipid class from compound properties derivable from MS1 spectra, namely their elemental molecular formula, can be stated as a supervised machine learning problem. With supervised machine learning, models are trained that predict a ‘label’ (e.g. lipid class) from a set of features (a feature vector) describing an input (e.g. elemental components of a molecular formula). These models are not constructed by hand, but rather, example inputs with known labels are used to ‘train’ a model. Using a large lipid database such as LIPIDMAPS (Sud *et al.*, 2007) that contains many examples of known lipids and their associated lipid classification, a set of generalized predictive models can be constructed via training to infer rules for predicting the correct lipid classes from elemental molecular formulae.

### A Chemically-Descriptive Feature Vector and an Appropriate Machine Learning Algorithm for Lipid Classification

The selection of both a chemically-descriptive feature vector and an appropriate machine learning algorithm will heavily influence both the performance and applicability of the resultant lipid class predictive models. Feature vectors must be sufficiently descriptive so that the algorithm has sufficient information to differentiate between inputs with different lipid class labels, but must also contain information that can be readily and accurately acquired through direct infusion FT-MS MS1 experiments. As such, structural information, which is the most informative, cannot be included in our feature vectors. Limiting our feature vectors to only information that can be acquired routinely from MS1, namely elemental molecular formula assignments provided by our SMIRFE algorithm, still provides substantial chemical information, including, the number of atoms for each element present in the formula and the theoretical monoisotopic mass.

Random Forest (Breiman, 2001) has been successfully applied to many metabolomics problems (Chen *et al.*, 2013) (Wang-Sattler *et al.*, 2012) and has several properties, which makes it an ideal machine learning method for this use-case. First, the classification of inputs into lipid classes is a highly hierarchical problem, for which Random Forest provides excellent performance. Second, a Random Forest of decision trees excels at learning classification rules based on discrete data like elemental atom counts. Third, the bagging process intrinsic to the Random Forest algorithm provides protection against overfitting and enables the direct measurement of classifier accuracy similar to explicit cross-validation (Svetnik *et al.*, 2003). Fourth, bagging and the construction of many independent binary classifiers makes unbalanced training datasets, where each label is not equally represented in the training data, less problematic. This last property is especially important, since a training dataset based on LIPIDMAPS will be highly unbalanced with respect to the different lipid classes (see Table 1).

**Table 1A:** LMSD + HMDB_non_Lipid Model Performance (Category) The accuracy and precision of category-level models trained on the LMSD + HMDB_non_lipid dataset demonstrates excellent accuracy on all classes and excellent precision for all classes apart from polyketides (76.7%). The polyketides represent a very diverse set of structures compared to other lipid classes which explains this discrepancy. The number of examples of each category highlights the unbalanced nature of this dataset and motivated the use of Random Forest for these models. Each model was trained as a one-class against all model (i.e. the Fatty Acyl [FA] model was trained using the set of known Fatty Acyls as true positives and all other examples as true negatives). Inclusion of the LMISSD provided no additional examples of Fatty Acyls, Polyketides, Saccharolipids, or Sterol Lipids and had minimal effect on the precision and accuracy of the models.

| **LMSD + HMDB_non_Lipid Model Performance (Category)** |  |  |  |
| --- | --- | --- | --- |
| **Category** | **Precision** | **Out of Bag Accuracy** | **Number of Examples** |
| Fatty Acyls [FA] | 0.841 | 0.901 | 2031 |
| Glycerolipids [GL] | 0.996 | 0.995 | 532 |
| Glycerophospholipids [GP] | 0.995 | 0.996 | 1886 |
| Polyketides [PK] | 0.767 | 0.885 | 1376 |
| Prenol Lipids [PR] | 0.989 | 0.971 | 473 |
| Saccharolipids [SL] | 1.000 | 0.998 | 102 |
| Sphingolipids [SP] | 0.999 | 0.993 | 1404 |
| Sterol Lipids [ST] | 0.934 | 0.972 | 824 |
| not_lipid | 0.928 | 0.799 | 7587 |

**Table 1B:** LMSD + LMISSD + HMDB_non_Lipid Model Performance (Category)

| **LMSD + LMISSD + HMDB_non_Lipid Model Performance (Category)** |  |  |  |
| --- | --- | --- | --- |
| **Category** | **Precision** | **Out of Bag Accuracy** | **Number of Examples** |
| Fatty Acyls [FA] | 0.838 | 0.939 | 2031 |
| Glycerolipids [GL] | 0.996 | 0.993 | 2715 |
| Glycerophospholipids [GP] | 0.979 | 0.980 | 9766 |
| Polyketides [PK] | 0.773 | 0.934 | 1376 |
| Prenol Lipids [PR] | 0.989 | 0.983 | 473 |
| Saccharolipids [SL] | 1.000 | 0.999 | 102 |
| Sphingolipids [SP] | 0.976 | 0.976 | 3089 |
| Sterol Lipids [ST] | 0.931 | 0.983 | 824 |
| not_lipid | 0.930 | 0.882 | 7587 |

## Materials and Methods

### Structure of Chemically-Descriptive Feature Vectors

As illustrated in Figure 1, the feature vector, based on a given molecular formula, contains an atom count for each CHONPS element, the sum of atom counts for other elements, and the theoretical monoisotopic mass and individual decimal places from this mass. To ensure that all molecular weights for all entries were comparable, every entry had its theoretical monoisotopic molecular mass re-calculated using isotope molecular masses from NIST (Wieser *et al.*, 2013) (Berglund and Wieser, 2011). Each element atom count is an integer, but for different elements the expected atom count range can vary significantly. For biological lipids in general, up to 300 hydrogen atoms could be expected, but only a few sulfur or phosphorous atoms are expected. The theoretical monoisotopic mass is a floating-point number between zero and a few thousand daltons, while each digit will be represented as an integer between 0 and 9. As a result, each feature in our feature vector will be on a different scale. Although these could be normalized to remedy the differences in scale, which is a requirement for some machine learning algorithms, the Random Forest algorithm does not have this limitation.

**Figure 1:**
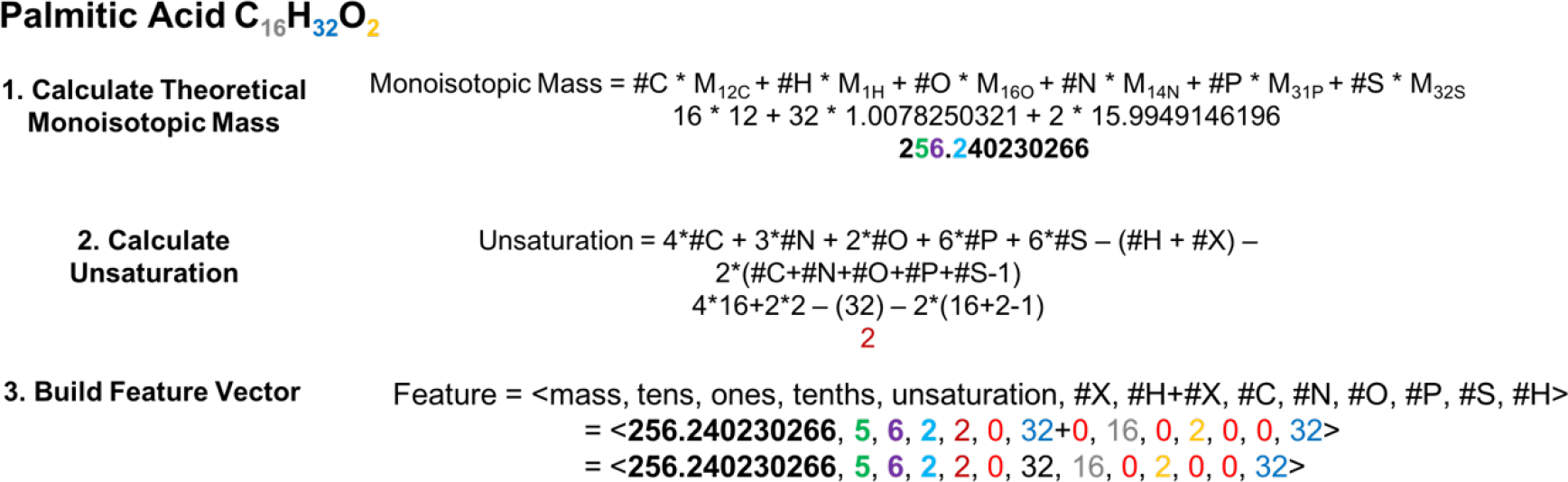
Example Feature Vector. Example construction of a feature vector for the EMF C_16_H_32_O_2_, corresponding to palmitic acid. In a real-world application, the EMF would be provided from an assignment method such as SMIRFE and the compound it represents may not be known. The first step in constructing the feature vector is to calculate the theoretical monoisotopic mass for that EMF. Calculating the theoretical mass for an EMF rather than relying upon the observed mass for the corresponding spectral feature, eliminates the potential confound of mass error at the classification step. Calculating and using the monoisotopic mass is necessary so that isotopologues of the same EMF can be classified using the same classifiers. In the second step, the number of hydrogens missing in the formula due to unsaturation is calculated. Finally, the monoisotopic mass, the number of missing hydrogens and the EMF are used to construct the feature vector. The coloring and bolding of the numbers in the example feature vector reflect the sources of these values.

**Figure 2:**
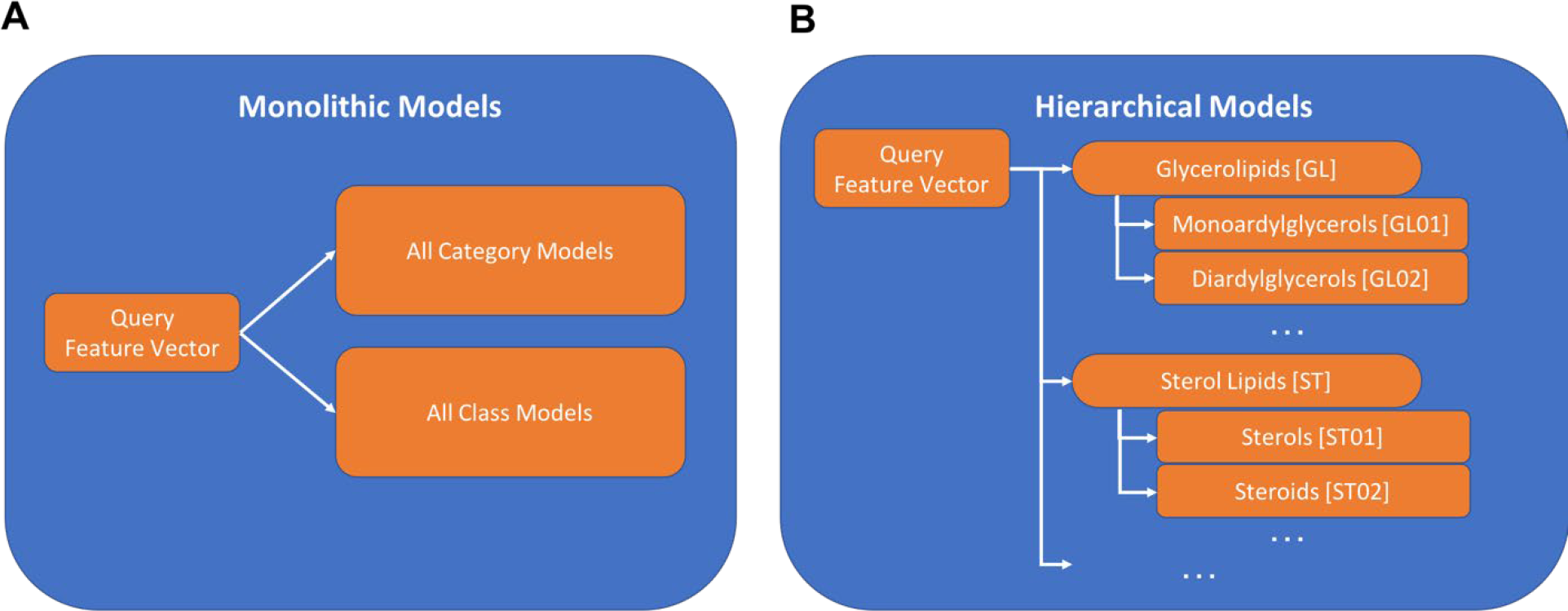
Organization of Hierarchical and Monolithic Models. In a monolithic organization there exists one model for classifying feature vectors into lipid categories and another model for lipid classes (Panel A). This organization is simpler with fewer model to train compared to the hierarchical organization of models (Panel B). In the hierarchical organization there are more models total but the class models are organized under their respective category model.

**Figure 3:**
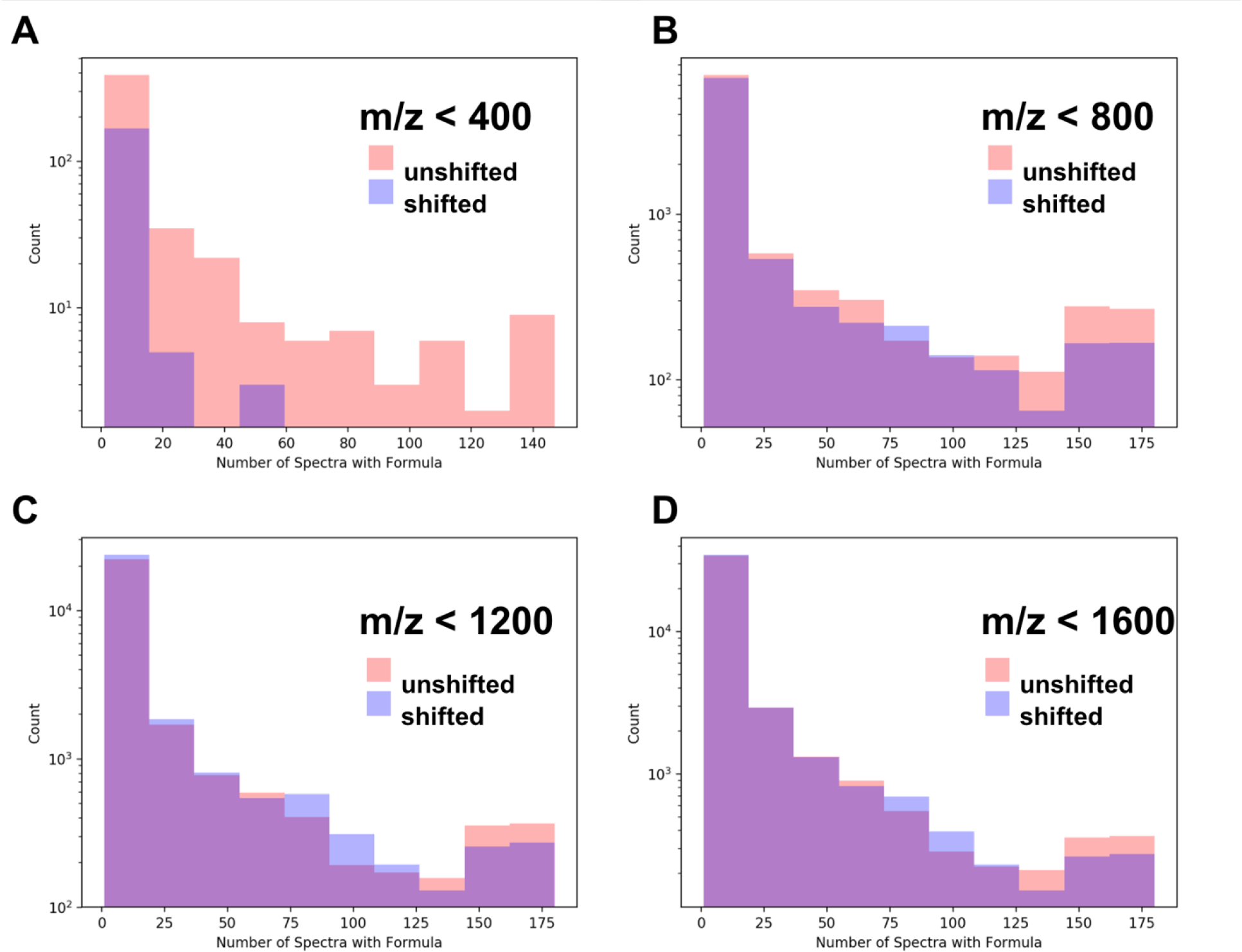
Cross-Sample Correspondence Identifies High Quality Assignments. Correct assignments are expected to occur more consistently within a set of samples than incorrect assignments. As shown in Panel A, below 400 m/z, very few assignments are made in the shifted spectra and very few of the assignments correspond across an appreciable number of spectra (i.e. the vast majority of the first bin represents single spectra assignments). As m/z increases (Panels B-D), shifted spectra have more assignments and by chance some of these assignments correspond in multiple samples. However, at up to 1200 m/z, there are clearly more well corresponding formulas in the unshifted assignments than in the shifted assignments. These results imply that assignment correspondence can be used to select correct assignments up to a given mass cutoff. In this dataset, this appears to be less than 1200 m/z.

### Derivation and Organization of Training Datasets

In addition to the selection of proper feature vectors and the selection of an appropriate machine learning algorithm, the quality of a machine learning model depends heavily on the quality of the training data from which the model is constructed. Training datasets should be large, contain examples of both true positives and true negatives, and cover most of the expected feature space. Additionally, training data must be organized in the appropriate manner. In this case, the training data should have the training inputs mapped to both high-level lipid categories (e.g. glycerolipid, phospholipid, etc.) and further subdivided into more specific “main classes” (e.g. monoradylglycerols, eicosanoids, secosteroids, etc.).

The LIPIDMAPS database is the largest lipid-specific repository of metabolite structures and every entry in LIPIDMAPS is assigned to both a high-level lipid category and a lower-level ee. There are 7 lipid categories, which are further subdivided into 79 distinct classes. Each entry represents either an observed lipid or a predicted lipid and contains an elemental formula for that lipid and its assigned lipid category and lipid class. Therefore, entries from LIPID MAPS are sources of true positives for our training dataset. LIPID MAPS is also subdivided into two databases: the LIPID MAPS Structure Database (LMSD) and the LIPID MAPS In-Silico Structure Database (LMISSD). The LMSD contains both manually verified and computationally generated lipids and is freely available for download, while the LMISSD is completely computationally generated lipids. Unlike the LMSD, the LMISSD is not directly downloadable and a web scraper written in R (Ihaka and Gentleman, 1996) using the RSelenium package (Harrison, 2016) was used to extract every LMISSD entry with its lipid category, lipid class, and molecular formula. We downloaded the LMSD in September, 2018, which contained 42,004 entries. We webscraped the LMISSD in September, 2018, obtaining 1,131,106 entries.

However, true positives are only one half of a training dataset. True negatives are also needed for the construction of robust models. In this case, a true negative is a biological formula that is not a lipid. The human metabolome database (HMDB) (Wishart *et al.*, 2012) contains many examples of biological formulas of known class and is freely downloadable. By filtering out and removing known lipids from the HMDB, a set of false negatives were constructed. These entries, of course, do not have a lipid category or lipid class assigned to them, so an extra category and class called ‘non_lipid’ was assigned to these entries. We downloaded version 4.0 of the HMDB on September, 2018, which contained 114,089 entries with 22,657 entries being non-lipids.

Since in-silico generated lipids may not necessarily exist in biological systems, it is prudent to construct two example training datasets: HMDB non_lipids + LMSD (referred to as LMSD training set) and HMDB non_lipids + LMSD + LMISSD (referred to as LMISSD training set). Since isomers of lipids can have the same molecular formula but have a different structure that can even belong to different lipid categories and lipid classes, each training dataset was deduplicated by mapping each formula to all observed lipid categories and classes for each formula. A large portion of the entries in both the LMSD and LMISSD are isomers of other entries of the same lipid class and category. The final LMSD + HMDB non_lipid training dataset resulted in 16,215 unique entries as compared to 30,692 for the LMSD + LMISSD + HMDB non_lipid training dataset.

### HMDB-Derived Molecular Formula Convex Hull Construction

From the set of HMDB formulas composed only of CHONPS elements and with a molecular weight below 1600 m/z, a convex hull was constructed and enumerated to generate theoretical metabolite formulae of biological origin. In this formulation, each molecular formula from the HMDB represents a point in a six-dimensional space (each dimension representing the number of a CHONPS element present in the formula) where each point has integer coordinates corresponding to the number of each element present. The convex hull around these points was constructed using the Python implementation of the qhull algorithm (Barber and Huhdanpaa, 1995). All possible points within the convex shape were then enumerated to generate all CHONPS-specific molecular formulae within the convex hull.

### Experimentally-Derived Molecular Formulae from Human Lung Cancer Samples

Paired cancer and non-cancer tissue samples were acquired from eighty-six patients with suspected resectable stage I or IIa primary non-small cell lung cancer (NSCLC). Specimens were obtained primarily using wedge resection and all specimens were harvested within 5 minutes after pulmonary vein clamping to minimize ischemia in the resected tissues. Immediately after resection, the tumor was transected and sections of cancerous tissue and surrounding non-cancer tissue at least 5 cm away from the tumor were immediately flash frozen in liquid nitrogen and stored at <80°C. On-site pathologists confirmed the diagnosis and cancer-free margins on parallel tissue samples. All samples were collected under a University of Louisville approved Internal Review Board protocol and written informed consent was obtained from all subjects prior to inclusion in the study. The frozen samples were then prepared and analyzed using two Thermo Orbitrap Fusion instruments interfaced to an Advion Nanomate nanoelectrospray source. Additional details on sample preparation and mass spectrometric analysis are included in supplemental materials.

MS1 spectra were acquired for each sample using direct infusion. These MS1 spectra were then assigned using our in-house SMIRFE assignment tool which assigns spectral features without a database of expected molecular formulas corresponding to metabolites. Instead, SMIRFE generates an exhaustive list of expected molecular formulas which can be queried using a peak’s observed m/z with a mass tolerance determined by the digital resolution of the instrument, which is approximately 1ppm for the Fusion instrument. SMIRFE uses patterns in the intensity ratios of suspected isotopologues of the same elemental molecular formula and how these patterns compare to predicted intensity ratios based on isotope natural abundances. Assignments were generated for 192 samples up to 1600 m/z and SMIRFE assigned 127,338 unique formulas.

### Monolithic Classifier Construction

Initially a single, monolithic random forest classifier was constructed for the simultaneous classification of all lipid categories and classes. This construction was done using the random forest implementation from sklearn (Pedregosa *et al.*, 2011) with default hyperparameters except for the number of decision trees, which were varied from the default of 10 trees to 500 trees.

### Hierarchical Classifier Construction and Organization

Using the monolithic organization, a single model exists for lipid categories and a second model exists for lipid classes (Panel A). Each query feature vector is processed by both models to produce category and class labels and each model can be used independently. While conceptually simple and easier to implement, the monolithic organization suffers from relatively poor performance.

An alternative approach is a hierarchy of models in which category and class models are combined (Panel B). Each class and category model is its own Random Forest model. This organization has several distinct advantages over the monolithic implementation. First, the hierarchal organization enables the simplification of each decision boundary that each model must learn and each model can select its optimal set of features for drawing that boundary. Second, using category classifiers to filter what feature vectors should be passed to lower level class models effectively results in machine learning models feeding their results into other machine learning models. This technique is employed in deep learning to construct robust and powerful classifiers for complicated classification problems. Third and finally, collections of relatively weak classifiers working together often outperform monolithic classifiers. This observation is also well-known in the machine learning field (Polikar, 2012) and is the central motivating concept behind ensemble machine learning algorithms such as Random Forest.

These advantages come at the cost of additional manual overhead to segment and organize the training datasets appropriately and additional computational overhead to construct and train multiple models. This cost is largely mitigated by the fact that models need to be trained only once (or very rarely) and can then be reused, effectively amortizing the cost of building the models and using preconstructed models for classification is computationally cheap. For all models, the Random Forest implementation from the Python sklearn package (Pedregosa *et al.*, 2011) was used with default hyperparameters except for the number of trees which was set to 500.

### Evaluation of Lipid Classification Performance

The performance of any machine learning model can be evaluated using a variety of metrics; however, rarely does a single metric fully reflect the goodness of any model in all use cases. For example, classifiers with high overall accuracy for the whole training set may classify certain labels very poorly which may not be obvious from a global accuracy metric. This is especially true for unbalanced training datasets, where conservative models will have high accuracy but make very few classifications. For this use-case, the chemical characterization of direct infusion MS1-based lipidomics experiments, models that perform well for all lipid labels are obviously desirable, but more importantly, high specificity is desired. Models that generate many incorrect assignments will, at the very least, become burdensome to utilize effectively, and at worst, could lead to incorrect interpretation of results. Therefore, highly accurate and highly precise models are desirable even at the cost of missing some true positive classifications.

To evaluate the accuracy for the Random Forest models, we used the out-of-bag training accuracy. While not strictly equivalent to explicit cross-validation, the accuracy metric provided by out-of-bag training accuracy is a sufficient proxy, even though for unbalanced datasets such as ours, the out-of-bag training accuracy *underestimates* the error rate (Janitza and Hornung, 2018); however, from our own experience, this underestimation is not significant. For precision, the classic definition of precision can be applied for sets of inputs on known label and their model-generated labeling. Models that are both highly accurate and highly precise are unlikely to generate false assignments and are suitable for our task.

## Results

### Monolithic Classifier Performance on Training Datasets

Using the LMSD + HMDB_non_lipid dataset, the performance of a monolithic classifier for lipid category and lipid class was tested. Even with 500 trees, the monolithic Random Forest models were only able to achieve an out-of-bag accuracy of 74.9% for lipid category and 87.3% for lipid class. Including the LMISSD resulted in an out-of-bag accuracy of 83.1% for lipid categories and 80.1% for lipid class. In both datasets, the presence of a large number of non-lipid entries inflates the lipid class accuracy as all non-lipid entries map to the non-lipid class. Although monolithic classifiers may have the theoretical advantage of being simpler to implement, train, and deploy, their usefulness is limited by their relatively poor classification performance.

### Multi-Classifier Performance on Training Datasets

For both training datasets (LMSD + HMDB_non_lipid and LMSD + LMISSD + HMDB_non_lipid), the out-of-bag accuracy and precision for each lipid category are shown in Table 1A, while the class level results are shown in Supplemental Table 1. For all categories, the LMSD + HMDB_non_lipid trained models achieved high precision and high accuracy for all lipid categories. Classification performance for lipid class varies between classes but is in general excellent for classes with enough examples. The LMISSD-trained models achieved similar precision and accuracy for all categories (Table 1B) and classes (Supplemental Table 2). Although individually high accuracy or high precision would not necessarily indicate a well-trained model, the combination of high accuracy and precision across the models implies that the combined classification performance is robust and can be effectively applied to experimentally-derived molecular formulae.

**Table 2A:** LMSD + HMDB_non_lipid Model Performance for Convex Hull (Category) The formulas within the convex hull surrounded by the HMDB metabolites represent a very large set of plausible metabolites formulas. Lipid categories were predicted for every formula within the hull. For all categories, more formulas were predicted for each category than existed in the training dataset, indicating that the models have generalized beyond the training dataset. The extent of this generalization varied depending on the training dataset. For example, saccharolipids (the category with the smallest number of examples in the training dataset) was predicted more frequently in the LMSD trained models than in the LMISSD models, while sphingolipids were more frequently predicted in the LMISSD trained models than in the LMSD trained models. Although the distribution of predicted lipid categories varies slightly between the two sets of models, the overall trends are comparable. For example, sphingolipids were the highest predicted lipid category from the convex hull dataset by both sets of models.

| **LMSD + HMDB_non_lipid Model Performance for Convex Hull (Category)** |  |  |
| --- | --- | --- |
| **Category** | **Predictions** | **% of Hull Formulas** |
| Fatty Acyls [FA] | 475,516 | 0.429 |
| Glycerolipids [GL] | 8,205 | 0.007 |
| Glycerophospholipids [GP] | 1,145,418 | 1.033 |
| Polyketides [PK] | 84,333 | 0.076 |
| Prenol Lipids [PR] | 18,684 | 0.016 |
| Saccharolipids [SL] | 6,708 | 0.006 |
| Sphingolipids [SP] | 7,494,579 | 6.761 |
| Sterol Lipids [ST] | 18,643 | 0.017 |
| not_lipid | 74,621,680 | 67.31 |
| no category | 29,202,459 | 26.34 |

**Table 2B:** LMSD + LMISSD +HMDB_non_lipid Model Performance for Convex Hull (Category)

| **LMSD + LMISSD +HMDB_non_lipid Model Performance for Convex Hull (Category)** |  |  |
| --- | --- | --- |
| **Category** | **Predictions** | **% of Hull Formulas** |
| Fatty Acyls [FA] | 393,314 | 0.354 |
| Glycerolipids [GL] | 56,116 | 0.051 |
| Glycerophospholipids [GP] | 1,735,925 | 1.566 |
| Polyketides [PK] | 118,968 | 0.107 |
| Prenol Lipids [PR] | 15,881 | 0.014 |
| Saccharolipids [SL] | 2,795 | 0.002 |
| Sphingolipids [SP] | 12,568,226 | 11.34 |
| Sterol Lipids [ST] | 15,670 | 0.014 |
| not_lipid | 73,562,707 | 66.36 |
| no category | 27808607 | 25.08 |

### Multi-Classifier Performance on Theoretical Molecular Formulae

Brute force enumeration and testing of all points within the convex hull constructed around all CHONPS-only molecular formulae in the HMDB identified 110,857,519 formulae. While a brute force approach was computationally expensive, requiring several thousand CPU-core hours of time, it was necessary due to memory constraints with more complex methods. However, classification of the resulting convex hull took approximately 10 CPU-core hours. Calculations were performed on a quad-socket system with four E7-4820v4 CPUs (10 cores, 20 threads each) clocked at 2.00 Ghz and 1TB of RAM clocked at 2400MHz. Classifying these formulae with the LMSD and LMISSD models resulted in the majority of formulas assigned to either the non_lipid category or to no category at all. Results for each category are summarized in Table 2. The LMISSD models predict 4 of the 7 categories more frequently than the LMSD models but the trend in predicted categories were similar. Given the number of formulas in the convex hull that do not correspond to ‘real’ metabolite formulas, a high percentage of non_lipid or no classification formulas is expected if our models are highly discriminating.

### Multi-Classifier Performance on Experimentally-Observed Molecular Formulae

The distribution of the assigned lipid categories on molecular formulae enumerated from a human lung cancer FT-MS dataset is shown in Table 3A for the LMSD based classifier. SMIRFE generates many possible assignments for each peak at higher m/z as the number of possible formulas increases dramatically with increasing m/z. As a result, a relatively small percentage of formulae are assigned to a lipid category but many peaks have at least one formula that was assigned to a lipid category. For the LMSD models, the ability to predict lipid category and class drops substantially after about 1200 m/z. This is due to the low number of entries in the LMSD at higher m/z.

**Table 3A:** LMSD + HMDB_non_lipid Model Performance for Unshifted Assignments. SMIRFE assignments were generated for the NSCLC dataset described in supplemental material. SMIRFE assignments are generated in an untargeted manner without using a database of known lipids. For the peak masses across all peaklists, 127,338 total formulas were assigned and then classified. 32,688 total lipid category classifications were made with the most commonly assigned categories being glycerophospholipids and sphingolipids. A similar result was observed in the convex hull results as well, potentially indicating that this is an artifact of the classification method or possibly that these lipid categories are much more diverse than other categories. When each peak was shifted by 21 m/z, roughly 3% more formulas were assigned and 6% more lipids were classified. This small relative increase in SMIRFE assignments is likely due to the increased search space density with a 21 m/z shift. More importantly, the large number of artifactual assignments reflects the necessity of high-quality data prior to classification. Methods that can predict high-quality assignments correctly are not necessarily protected from the effects of low-quality spectral data that can cause misassignment.

| **LMSD + HMDB_non_lipid Model Performance for Unshifted Assignments** |  |  |
| --- | --- | --- |
| **Category** | **Predictions** | **% of Assigned Formulas** |
| Fatty Acyls [FA] | **639** | 0.502 |
| Glycerolipids [GL] | 795 | 0.624 |
| Glycerophospholipids [GP] | 8062 | 6.331 |
| Polyketides [PK] | 28 | 0.022 |
| Prenol Lipids [PR] | 1054 | 0.827 |
| Saccharolipids [SL] | 166 | 0.130 |
| Sphingolipids [SP] | 21586 | 16.952 |
| Sterol Lipids [ST] | 358 | 0.281 |
| not_lipid | 54389 | 42.71 |
| no category | 40683 | 31.95 |

**Table 3B:** LMSD + HMDB_non_lipid Model Performance for Shifted Assignments.

| **LMSD + HMDB_non_lipid Model Performance for Shifted Assignments** |  |  |
| --- | --- | --- |
| **Category** | **Predictions** | **% of Assigned Formulas** |
| Fatty Acyls [FA] | **258** | 0.1951 |
| Glycerolipids [GL] | 923 | 0.7001 |
| Glycerophospholipids [GP] | 9517 | 7.227 |
| Polyketides [PK] | 37 | 0.0281 |
| Prenol Lipids [PR] | 1160 | 0.8808 |
| Saccharolipids [SL] | 233 | 0.1769 |
| Sphingolipids [SP] | 22370 | 16.99 |
| Sterol Lipids [ST] | 257 | 0.1952 |
| not_lipid | 51863 | 39.38 |
| no category | 45663 | 34.67 |

When the masses of the peaks are shifted by +21 m/z to mimic a gross miscalibration error, the number of SMIRFE assignments is increased, from 127,338 to 131,690 total formulas and the number of predicted lipids increases as well from 32,688 to 34,755 (Tables 3A and 3B). This result implies that the lipid classifier cannot be used alone to screen out all bad assignments when lipids are expected, instead other orthogonal data must be used to verify the quality of the assignments and select the correct assignments.

### Cross-Sample Assignment Correspondence Improves Assignment Quality

Limits in mass resolution and intensity resolution and the immense size of the search space considered by SMIRFE at high masses leads to ambiguous assignments for many peaks. When mass error is present, ambiguous and incorrect assignments can be generated. However, the correct assignment for a peak should be assigned more consistently for a consistently observed feature in the dataset. Therefore, how well an assignment corresponds across samples in a dataset is a potential avenue for selecting high quality assignments. Figure C shows histograms of assignment correspondence for elemental molecular formulas derived the spectra of the lung cancer dataset. Much higher correspondence is observed in the unshifted assignments vs the shifted assignments and the number of shifted high-correspondence assignments are fewer at lower m/z.

## Discussion

### Classifier Organization and Performance

As mentioned previously, a monolithic, multi-lipid-class predictive model failed to achieve top performance for the task of classifying assigned molecular formulas into lipid categories and classes. We hypothesize that this is due to the inability of a single classifier to represent all these boundaries completely and accurately. This single classifier must not only learn how to separate lipids from non-lipids, but it must also subdivide the lipid feature space into discrete spaces representing each category and further subdivide these category spaces into class spaces. Much of this subdivision can be done explicitly during training. For example, the diacylglycerols are a sub-class of the larger category of glycerolipids and a less powerful classifier can easily identify the diacylglycerols from other glycerolipids when it must only learn that single decision boundary. As a result, our organization of weaker predictive models had superior performance. Initially, this behavior can seem counterintuitive but is consistent with the concept of ensemble learning from machine learning where collections of weaker classifier models often outperform fewer larger classifier models when properly organized. The hierarchy of models that are constructed mirror how a human would approach the classification problem. For example, if a molecular formula is known not to be of the sphingolipid category, a human will not attempt to assign this formula to a sphingolipid class; however, a monolithic model will attempt to do so. This wastes computational power and increases the likelihood of incorrect prediction of both class and category.

The final models produced by our tool achieved both high accuracy and high sensitivity on the training dataset. Of course, performance on training data does not paint a complete picture of model performance, but for Random Forest which implements bagging, these metrics predict performance on inputs similar to the training data. Models with both high accuracy and high sensitivity are unlikely to produce incorrect lipid assignments, but may be overly conservative and fail to generate a non_lipid assignment for some inputs. While this behavior is undesirable, it is preferable to less conservative models that will yield many incorrect lipid category and class predictions.

### LMSD vs LMISSD Trained Models

One method for improving the performance of a machine learning model is to provide larger amounts of training data, which in turn enables more informed and more accurate decision boundaries to be determined. For this reason, models were trained using both the LMSD and LMISSD, which has nearly 25 times the number of entries as the LMSD. However, LMISSD trained models did not offer substantially improved performance as compared to the LMSD-only models on the training datasets. Although the LMISSD contained many entries, the input training set only doubled in size after isomeric entries were removed, implying that little information was added regarding the distribution of formulas in lipid category or lipid class space. Another possible explanation for this observation is that the LMISSD contains substantially more entries, but for only 4 out of the 7 categories in the LIPIDMAPS database and that the decision boundaries for these categories were already well-determined by the LMSD only models.

### Classifier Generalization

A benefit that machine learning models have over traditional database lookups are their ability to infer rules that can be applied to never observed inputs to make accurate predictions. This ability was demonstrated with both the LMSD- and LMISSD-constructed models. Both models produced lipid category and class predictions for experimental and theoretical molecular formulae not present in the training dataset. However, the generalizability of the models depends heavily on the quality and size of the input dataset (Supplemental Figure 1).

Despite having similar performance on the training datasets, the LMISSD and LMSD trained models had similar but distinctive behavior on the convex hull metabolites. The LMISSD assigned many more sphingolipids than the LMSD models and in general, the categories with more examples in the training dataset were more frequently predicted. This could be due to a bias in the trained models from the unbalanced training data or could reflect the relative amount of structural diversity possible within each class, i.e. the number of possible sphingolipid formulas might truly be larger than the number of possible sterol formulas. However, the percentage of hull formulas predicted for each category was similar between the two models, implying that they are overall very similar. Both models predicted roughly the same number of non_lipid formulas implying that the overall lipid vs non-lipid decision boundaries of the two models are very similar. Discrepancies between the two models can also be attributed to the presence of predicted lipids in the LMISSD that do not exist – this could confuse classifiers if the predicted lipids and the validated lipids suggest different decision boundaries.

Ultimately the ability of both models to make accurate predictions will be improved with larger training datasets. With more examples that more exhaustively span lipid formula space, the more accurate and generalizable the models constructed using these same methods will become. However, given the marginal improvement with a doubling of the training dataset from the LMSD vs LMISSD, improvements may be marginal without a vastly larger training dataset.

### Mass Error and Classification Results

Ideally, a substantial mass error would result in no formulas being assigned by SMIRFE or that the assigned formulas fail to classify. As shown with our NSCLC dataset, a large mass error does not eliminate all assignments nor completely abolish our ability to classify the resulting, almost certainly incorrect, assigned formulas.

Given the very large search space that an untargeted tool must search to generate assignments, almost any m/z has many possible assignments, given the theoretical molecular formula search space. Since a systematic error does not change the mass difference between isotopologues, patterns of isotopologues for these incorrect formulas can still be identified and assigned. Thus, without extremely high mass resolution to restrict the set of possible assignments considerably, which still may not be effective (Kind and Fiehn, 2006), a constant mass error will still produce assignments. Furthermore, current variance in peak intensities is not low enough to prevent artifactual assignment at higher m/z.

As was seen in the convex hull analysis, approximately a quarter of the generated formulas appeared to be lipids to the models. This could reflect the true distribution of lipids in possible formula space, but more likely it represents limitations of our models. Nonsense formulas that can arise from m/z error or from the convex hull method cannot be properly learned as they are very different from the training set data. Although the ability of our models to produce no classification for an input feature vector protects against this effect, it is not perfect. The same models that learn real (biochemically relevant) metabolite formulas correctly may fail to properly handle nonsense formulas that SMIRFE can assign to peaks with high mass error and noise or artifactual peaks.

Therefore, lipid classification alone should not be used to filter out features in datasets, especially on a single-spectrum basis. Information such as how many times a formula is observed across a dataset appears to be useful for filtering. Also, observed correlation between features classified to the same lipid category and/or class should provide additional discriminating criteria. Features considered trustworthy by this information and other methods can then be used for further analysis. Similar problems exist with targeted assignment tools as well and a lack of substantial cross-sample formula correspondence is an indication that there is a possible data quality problem preventing accurate assignment.

### Implications for Experimental Design

The ability to predict lipid category and class from molecular formula assignments without the need for cross-validated *metabolite* assignments, enables simpler experimental designs as the volume of information needed to perform class or category level comparisons is lessened. As molecular formula can be assigned from direct infusion FT-MS MS1 spectra directly and in a cross-validating manner, chromatography and other cross-validation information is not necessary for class or category level comparisons when using these models. However, the quality of the analyses will depend heavily on the quality of the assigned molecular formulae.

SMIRFE leverages patterns in the relative heights of isotopologue peaks for the same elemental molecular formula to determine what molecular formulae best explain features observed in a spectrum. Although SMIRE is not necessarily limited to only high-end mass spectrometers such as FT-MS instruments, only these instruments provide enough mass accuracy and resolution to observe and characterize relevant sets of isotopologues. This restriction is becoming increasingly less relevant as high-performance spectrometers become more available. Additionally, SMIRFE and subsequent lipid prediction does not enable the robust assignment of metabolite structures to spectral features and this will still require additional information from orthogonal experiments.

### Conclusions

With untargeted analysis methods, lipidomics has the potential to produce more informative datasets that will aid in the construction of more complete models of cellular metabolism. This in turn enables a better understanding of both healthy and disease processes. A necessary step in many of these analyses is the assignment of lipid category or class to an observed lipid feature. When multiple orthogonal sources of information are available (i.e., MS + chromatography, NMR + chromatography, MS/MS), lipid category and class assignment can be inferred from trustworthy metabolite assignments based on comparison to spectral databases; however, this approach limits untargeted analysis, since spectral and lipid databases are incomplete.

The application of machine learning algorithms enables the construction of models that can accurately and precisely assign lipid labels to observed spectral features that have been assigned to a molecular formula. Unlike other approaches that leverage metabolite databases directly for lipid assignment, these models have the capacity to infer lipid category and class for entries not present in existing databases. This capacity is essential for untargeted metabolomics experiments as database incompleteness can lead to a biasing of lipid classification and in turn biological interpretation. Since these models are informed by the existing metabolite databases during training, their capacity to compensate for database incompleteness is not unlimited as observed with our LMSD informed models having limited efficacy at higher mass ranges. The inclusion of additional sources of empirically observed lipids in these mass ranges may extend the useful mass range of this methodology. LMISSD-informed models did not suffer from this limitation, but had decreased accuracy and specificity, potentially attributable to unrealistic entries in the LMISSD.

By recapitulating the same level of performance observed with the training datasets, the robust ability of these models to make lipid classifications on molecular formulas assigned to features derived from spectra of non-polar tissue extracts demonstrates their potential for real-world application. Thus, machine learning-based approaches will allow for more untargeted lipid profiling analyses than existing database-centric methods, even with the more limited data that can be acquired using direct injection MS1 alone. Similar methods could be applied to the classification of other major types of biomolecules or to identify potential contaminants or non-biological compounds detected in complex biological samples. However, the quality of the predictions made by such methods remains limited by the ability to generate high quality assignments in an unbiased manner for higher m/z ranges that are relevant to lipid profiling. Methods such as SMIRFE combined with cross-sample correspondence provides a potential avenue to generating such assignments for these higher m/z ranges.

## Acknowledgements

This work was supported in part by grants NSF 1419282 (PI Moseley) and NIH UL1TR001998-01 (PI Kern). We thank Richard M. Higashi, Teresa W.-M. Fan and Andrew N. Lane who collected the human tissue samples. We thank Robert M. Flight for writing the web scraper for the LMISSD.

## Author Contributions

JMM designed and implemented the machine learning models and the convex hull analysis. HNBM designed the m/z shifted formula analysis and JMM implemented it. The primary manuscript writers were JMM and HNBM.

## Resource Sharing

Code and data used for this manuscript are available here: https://figshare.com/s/187eb261983b6d0aca1c

## Compliance with Ethical Standards

### Disclosure of potential conflicts of interest

The authors declare that they have no conflict of interest.

### Ethical Approval

The base study providing the de-identified data analyzed was approved by IRB protocol (IRB 14-0288-F6A) at the University of Kentucky and IRB protocol (#523.05) at the University of Louisville.

### Informed Consent

Written consent was obtained for the collection of human tissue samples under an IRB approved protocol (IRB 14-0288-F6A) at the University of Kentucky and IRB protocol (#523.05) at the University of Louisville.

## Supporting Information

These pages contain supporting information including descriptions of the tissue samples from which the experimental set of formulas was derived, additional result tables for the machine learning models, and a figure showing the distribution of molecular formulas across our training datasets with respect to m/z.

### Paired Human NSCLC Cancer and Non-Cancer Tissue Samples

Eighty-six patients with suspected resectable stage I or IIa primary non-small cell lung cancer (NSCLC) and without diagnosed diabetes were recruited based on their surgical eligibility. The extent of resection was determined by the surgeon in accordance with clinical criteria. Many of the specimens were obtained from wedge resections which minimizes surgery time while the other specimens were acquired in less than 5 minutes after the pulmonary vein was clamped. Both techniques minimize ischemia in the resected tissues. Immediately after resection, the tumor was transected and section of cancerous tissue and surrounding non-cancer tissue at least 5 cm away from the tumor were immediately flash frozen in liquid nitrogen and stored at <80°C. On-site pathologists confirmed the diagnosis and cancer-free margins on parallel tissue samples. All samples were collected under a University of Louisville approved Internal Review Board (IRB) protocol and written informed consent was obtained from all subjects prior to inclusion in the study (Sellers *et al.*, 2015).

The frozen samples were pulverized under liquid nitrogen to <10 μm particles using a Spex freezer mill, and extracted using a modified Folch method as previously described (Ren *et al.*, 2014). The lipid fraction was supplemented with 1 mM butylated hydroxytolune and then dried by vacuum centrifugation at room temperature. Samples for FT-MS analysis were redissolved 200-500 μl chloroform/methanol (2:1) supplemented with 1 mM butylated hydroxytolune. Reconsitituted lipids samples were diluted in in isopropanol/methanol/chloroform 4/2/1 (v/v/v) with 20 mM ammonium formate (95 μl of solvent for 5 μl of sample) before direct infusion.

### Mass Spectrometry Analysis of Tissue Samples

Ultrahigh resolution (UHR) mass spectrometry was carried out on a Thermo Orbitrap Fusion interfaced to an Advion Nanomate nanoelectrospray source using the Advion “type A” chip, also from Advion, inc. (chip p/n HD_A_384). The nanospray conditions on the Advion Nanomate were as follows: sample volume in wells in 96 well plate – 50 µl, sample volume taken up by tip for analysis – 15 µl, delivery time – 16 minutes, gas pressure – 0.4 psi, voltage applied – 1.5 kV, polarity – positive, pre-piercing depth – 10 mm. The Orbitrap Fusion Mass Spectrometer method duration was 15 minutes, and the MS conditions during the first 7 minutes were as follows: scan type – MS, detector type – Orbitrap, resolution – 450,000, lock mass with internal calibrant turned on, scan range (*m/z*) – 150-1600, S-Lens RF Level (%) – 60, AGC Target – 1e5, maximum injection time (ms) – 100, microscans – 10, data type – profile, polarity – positive. For the next 8 minutes, the conditions were as follows for the MS/MS analysis: MS properties: detector type – Orbitrap, resolution – 120,000, scan range (*m/z*) – 150-1600, AGC Target – 2e5, maximum injection time (ms) – 100, microscans – 2, data type – profile, polarity – negative; monoisotopic precursor selection – applied, top 500 most intense peaks evaluated with minimum intensity of 5e3 counts; data dependent MS^n^ scan properties: MS^n^ level – 2, isolation mode – quadrupole, isolation window (*m/z*) – 1, activation type – HCD, HCD collision energy (%) – 25, collision gas – Nitrogen, detector – Orbitrap, scan range mode – auto *m/z* normal, Orbitrap resolution – 120,000, first mass (*m/z*) – 120, maximum injection time (ms) – 500, AGC target – 5e4, data type – profile, polarity – positive. The ion transfer tube temperature was 275°C. (Yang *et al.*, 2017).

### Supplemental Tables 1 and 2

Supplemental Table 1 shows the training accuracy and precision for each class-level model in the model collection trained using the LMSD and the HMDB_non_lipid dataset. Excellent precision and accuracy were achieved for most classes; however, some classes have very few examples. For small classes, metrics of accuracy and precision are less useful. During class training, the category of each lipid is known – for the non-lipid category this results in perfect accuracy and precision, because there is only one non-lipid class for the non-lipid category. Each class is trained using a one-against all approach, which inflates the accuracy metric but not precision for each class model.

Supplemental Table 2 shows the same results on the LMSD + LMISSD + HMDB_non_lipid dataset. Similar metrics of precision and accuracy were achieved on this larger dataset.

**Supplemental Table 1.**
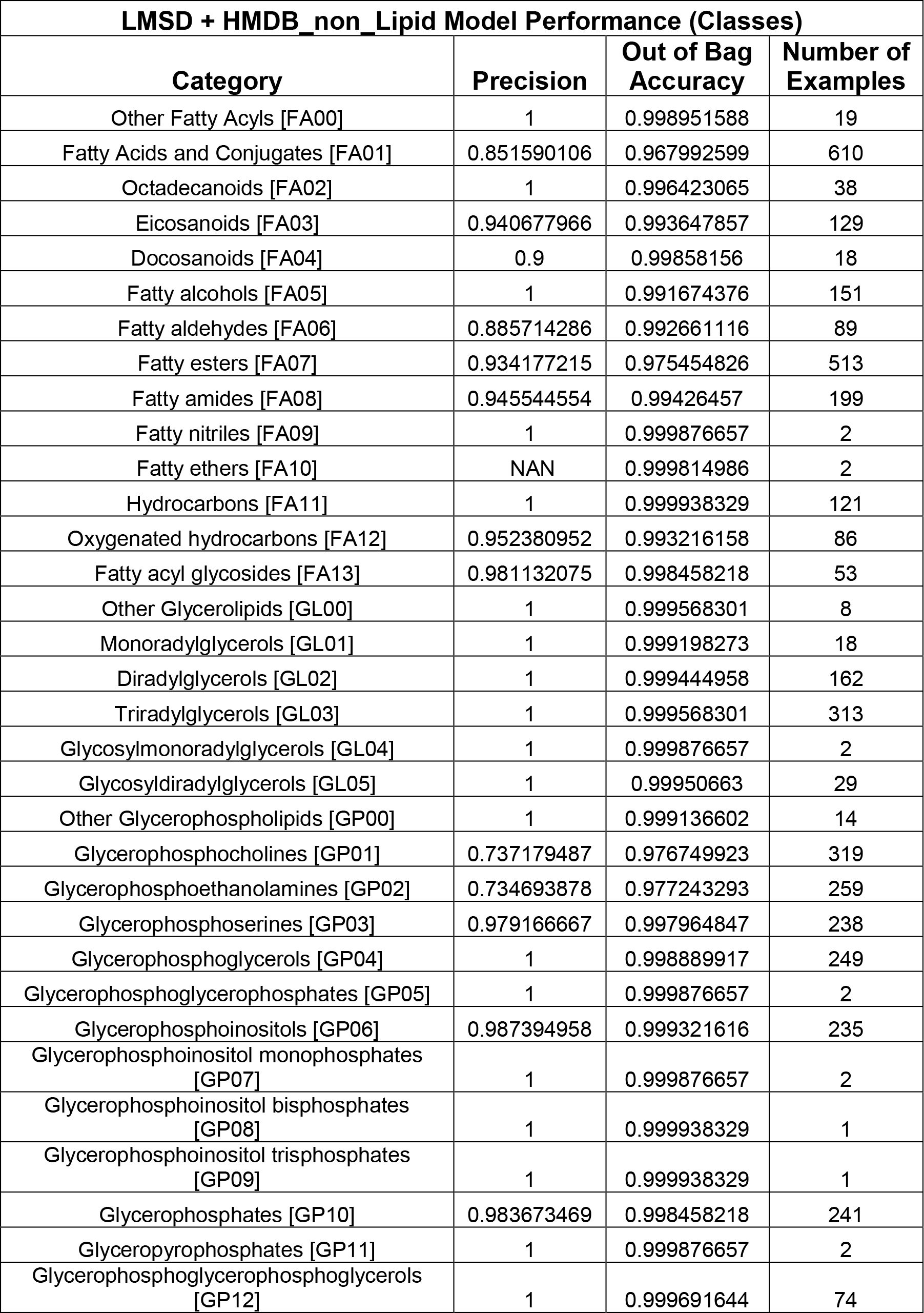
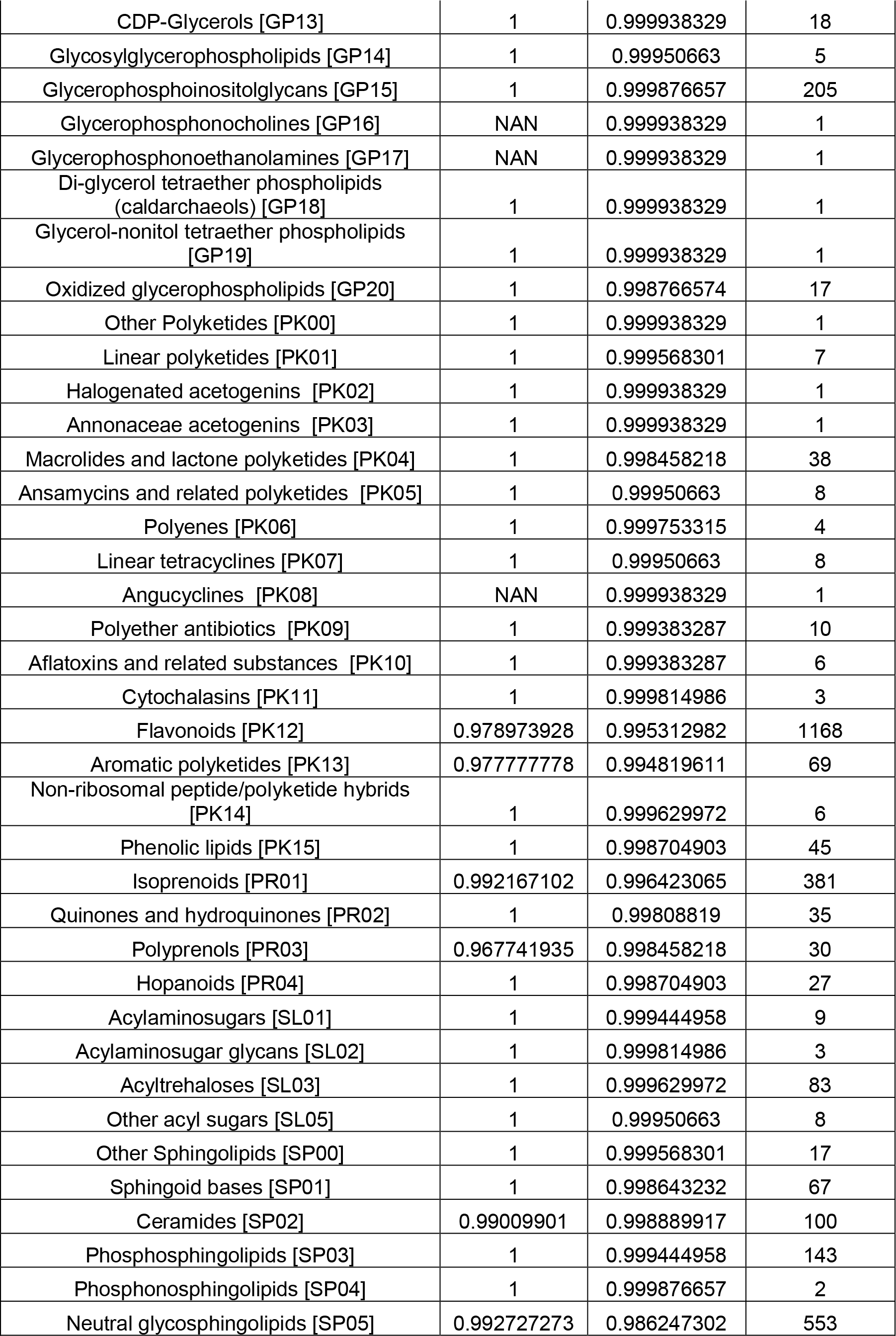
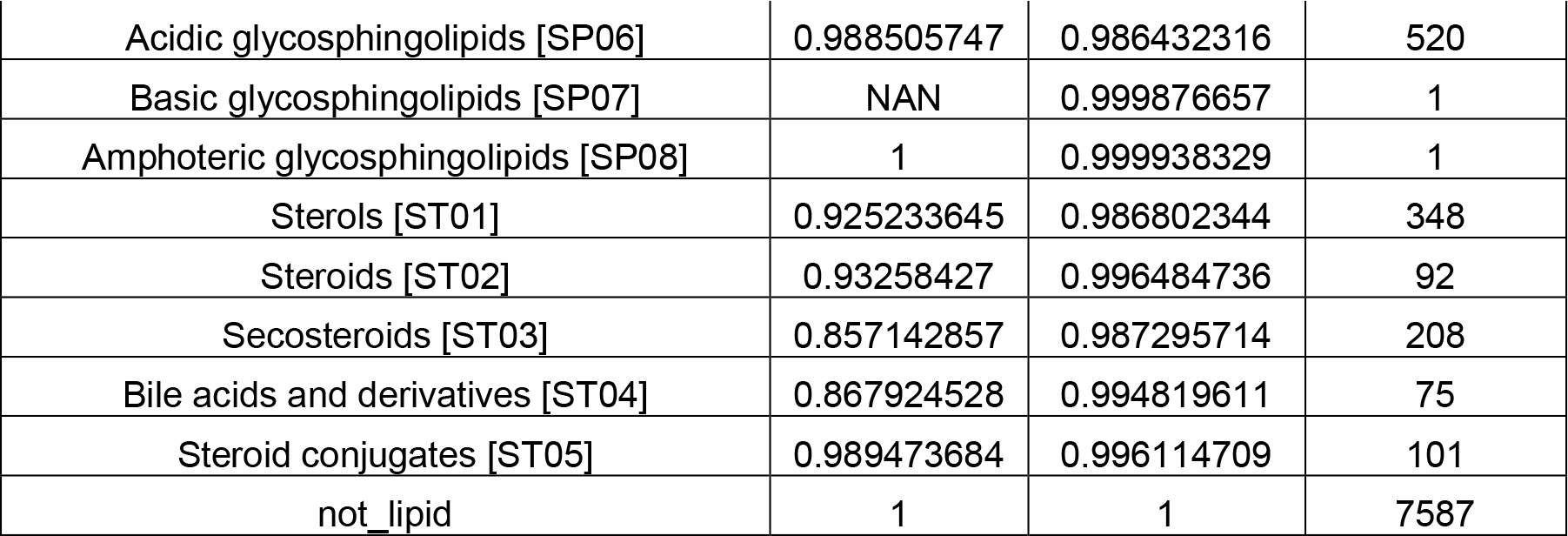

**Supplemental Table 2.**

| **LMSD + LMISSD + HMDB_non_Lipid Model Performance (Classes)** |  |  |  |
| --- | --- | --- | --- |
| **Category** | **Precision** | **Out of Bag Accuracy** | **Number of Examples** |
| Other Fatty Acyls [FA00] | 1 | 0.999392054 | 19 |
| Fatty Acids and Conjugates [FA01] | 0.850352113 | 0.980867575 | 610 |
| Octadecanoids [FA02] | 1 | 0.997890069 | 38 |
| Eicosanoids [FA03] | 0.964285714 | 0.9962808 | 129 |
| Docosanoids [FA04] | 0.833333333 | 0.999213246 | 18 |
| Fatty alcohols [FA05] | 1 | 0.995315238 | 151 |
| Fatty aldehydes [FA06] | 0.846153846 | 0.995672853 | 89 |
| Fatty esters [FA07] | 0.957333333 | 0.985945714 | 513 |
| Fatty amides [FA08] | 0.954545455 | 0.996638415 | 199 |
| Fatty nitriles [FA09] | 1 | 0.999928477 | 2 |
| Fatty ethers [FA10] | NAN | 0.999928477 | 2 |
| Hydrocarbons [FA11] | 1 | 0.999964238 | 121 |
| Oxygenated hydrocarbons [FA12] | 1 | 0.996101992 | 86 |
| Fatty acyl glycosides [FA13] | 0.963636364 | 0.9990702 | 53 |
| Other Glycerolipids [GL00] | 1 | 0.999713908 | 8 |
| Monoradylglycerols [GL01] | 1 | 0.998354969 | 61 |
| Diradylglycerols [GL02] | 0.813163482 | 0.984551014 | 492 |
| Triradylglycerols [GL03] | 0.915625 | 0.985945714 | 1242 |
| Glycosylmonoradylglycerols [GL04] | 1 | 0.999928477 | 2 |
| Glycosyldiradylglycerols [GL05] | 1 | 0.999713908 | 910 |
| Other Glycerophospholipids [GP00] | 1 | 0.999392054 | 14 |
| Glycerophosphocholines [GP01] | 0.719298246 | 0.968351035 | 571 |
| Glycerophosphoethanolamines [GP02] | 0.693939394 | 0.967850374 | 528 |
| Glycerophosphoserines [GP03] | 0.778012685 | 0.973214605 | 517 |
| Glycerophosphoglycerols [GP04] | 0.842436975 | 0.979758967 | 525 |
| Glycerophosphoglycerophosphates [GP05] | 1 | 0.999928477 | 512 |
| Glycerophosphoinositols [GP06] | 0.829787234 | 0.978221221 | 514 |
| Glycerophosphoinositol monophosphates [GP07] | 0.983050847 | 0.992382792 | 617 |
| Glycerophosphoinositol bisphosphates [GP08] | 0.911870504 | 0.9962808 | 509 |
| Glycerophosphoinositol trisphosphates [GP09] | 0.911870504 | 0.9962808 | 509 |
| Glycerophosphates [GP10] | 0.909090909 | 0.991345707 | 519 |
| Glyceropyrophosphates [GP11] | 1 | 0.999928477 | 2 |
| Glycerophosphoglycerophosphoglycerols [GP12] | 1 | 0.999749669 | 315 |
| CDP-Glycerols [GP13] | 1 | 0.999964238 | 18 |
| Glycosylglycerophospholipids [GP14] | 1 | 0.999713908 | 5 |
| Glycerophosphoinositolglycans [GP15] | 1 | 0.999785431 | 205 |
| Glycerophosphonocholines [GP16] | NAN | 0.999964238 | 1 |
| Glycerophosphonoethanolamines [GP17] | NAN | 0.999964238 | 1 |
| Di-glycerol tetraether phospholipids (caldarchaeols) [GP18] | 1 | 0.999964238 | 1 |
| Glycerol-nonitol tetraether phospholipids [GP19] | 1 | 0.999964238 | 1 |
| Oxidized glycerophospholipids [GP20] | 0.884877556 | 0.911132568 | 3882 |
| Other Polyketides [PK00] | 1 | 0.999964238 | 1 |
| Linear polyketides [PK01] | 1 | 0.999749669 | 7 |
| Halogenated acetogenins [PK02] | 1 | 0.999964238 | 1 |
| Annonaceae acetogenins [PK03] | 1 | 0.999964238 | 1 |
| Macrolides and lactone polyketides [PK04] | 1 | 0.998998677 | 38 |
| Ansamycins and related polyketides [PK05] | 1 | 0.999713908 | 8 |
| Polyenes [PK06] | 1 | 0.999856954 | 4 |
| Linear tetracyclines [PK07] | 1 | 0.999713908 | 8 |
| Angucyclines [PK08] | NAN | 0.999964238 | 1 |
| Polyether antibiotics [PK09] | 1 | 0.999642385 | 10 |
| Aflatoxins and related substances [PK10] | 1 | 0.999642385 | 6 |
| Cytochalasins [PK11] | 1 | 0.999892715 | 3 |
| Flavonoids [PK12] | 0.978973928 | 0.997282123 | 1168 |
| Aromatic polyketides [PK13] | 0.977777778 | 0.996924507 | 69 |
| Non-ribosomal peptide/polyketide hybrids [PK14] | 1 | 0.999785431 | 6 |
| Phenolic lipids [PK15] | 0.977777778 | 0.999105961 | 45 |
| Isoprenoids [PR01] | 0.98961039 | 0.997997354 | 381 |
| Quinones and hydroquinones [PR02] | 1 | 0.998962915 | 35 |
| Polyprenols [PR03] | 0.967741935 | 0.9990702 | 30 |
| Hopanoids [PR04] | 1 | 0.999284769 | 27 |
| Acylaminosugars [SL01] | 1 | 0.999678146 | 9 |
| Acylaminosugar glycans [SL02] | 1 | 0.999892715 | 3 |
| Acyltrehaloses [SL03] | 1 | 0.999713908 | 83 |
| Other acyl sugars [SL05] | 1 | 0.999713908 | 8 |
| Other Sphingolipids [SP00] | 1 | 0.999749669 | 17 |
| Sphingoid bases [SP01] | 1 | 0.999105961 | 67 |
| Ceramides [SP02] | 0.998548621 | 0.999678146 | 688 |
| Phosphosphingolipids [SP03] | 1 | 0.999785431 | 675 |
| Phosphonosphingolipids [SP04] | 1 | 0.999928477 | 2 |
| Neutral glycosphingolipids [SP05] | 0.996412556 | 0.992454315 | 1118 |
| Acidic glycosphingolipids [SP06] | 0.990384615 | 0.992239745 | 520 |
| Basic glycosphingolipids [SP07] | NAN | 0.999928477 | 1 |
| Amphoteric glycosphingolipids [SP08] | 1 | 0.999964238 | 1 |
| Sterols [ST01] | 0.927899687 | 0.992382792 | 348 |
| Steroids [ST02] | 0.93258427 | 0.997997354 | 92 |
| Secosteroids [ST03] | 0.886904762 | 0.99234703 | 208 |
| Bile acids and derivatives [ST04] | 0.897959184 | 0.996888746 | 75 |
| Steroid conjugates [ST05] | 0.979381443 | 0.997711261 | 101 |
| not_lipid | 1 | 1 | 7587 |

**Supplemental Figure 1:**
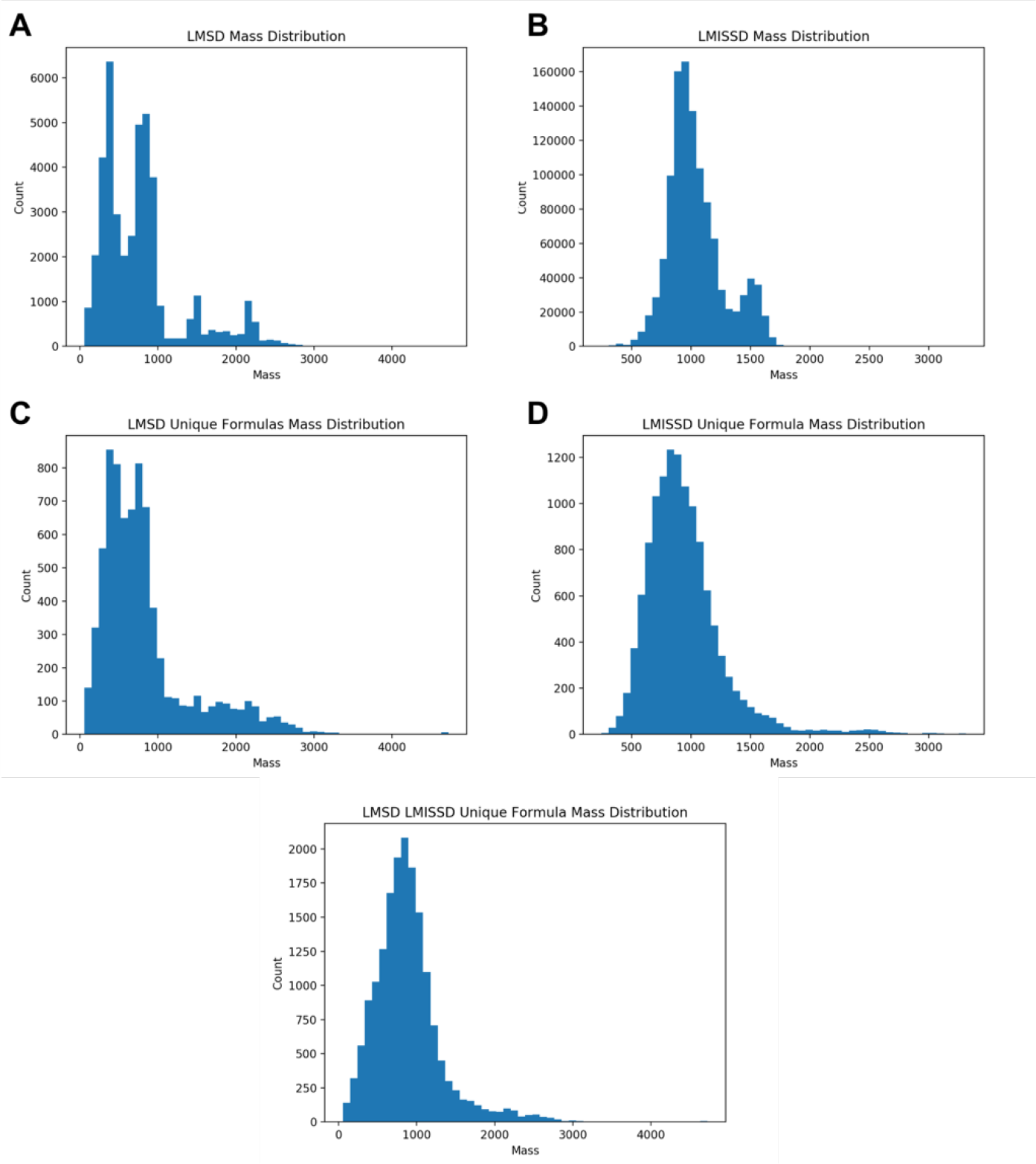
Training Set Mass Distributions. Both the LMSD and LMISSD are heavily biased towards lipids with a mass below 1200 m/z (Panels A and B respectively). This effect becomes clearer once entries are deduplicated to yield only unique formulas (Panels C and D). Some entries exist out to 3000+ m/z but the bulk of the formulas still reside in the sub 1200 mass range. When combined, the bias is still present in the combined set of unique formulas (Panel E). Additionally, the LMISSD does not have entries from all the lipid categories.

